# An optical aptamer-based cytokine nanosensor detects macrophage activation by bacterial toxins

**DOI:** 10.1101/2024.04.05.588290

**Authors:** Amelia K. Ryan, Syeda Rahman, Ryan M. Williams

## Abstract

Overactive or dysregulated cytokine expression is hallmark of many acute and chronic inflammatory diseases. This is true for acute or chronic infection, neurodegenerative diseases, autoimmune diseases, cardiovascular disease, cancer, and others. Cytokines such as interleukin-6 (IL-6) are known therapeutic targets and biomarkers for such inflammatory diseases. Platforms for cytokine detection are therefore desirable tools for both research and clinical applications. Single-walled carbon nanotubes (SWCNT) are versatile nanomaterials with near-infrared fluorescence that can serve as transducers for optical sensors. When functionalized with an analyte-specific recognition element, SWCNT emission may become sensitive and selective towards the desired target. SWCNT-aptamer sensors are easily assembled, inexpensive, and biocompatible. In this work, we introduced a nanosensor design based on SWCNT and a DNA aptamer specific to IL-6. We first evaluated several SWCNT-aptamer constructs based on this simple direct complexation method, wherein the aptamer both solubilizes the SWCNT and confers sensitivity to IL-6. The sensor limit of detection, 105 ng/mL, lies in the relevant range for pathological IL-6 levels. Upon investigation of sensor kinetics, we found rapid response within seconds of antigen addition which continued over the course of three hours. We found that this sensor construct is stable, and the aptamer is not displaced from the nanotube surface during IL-6 detection. Finally, we investigated the ability of this sensor construct to detect macrophage activation caused by bacterial lipopolysaccharides (LPS) in an in vitro model of disease, finding rapid and sensitive detection of macrophage-expressed IL-6. We are confident further development of this sensor will have novel implications for diagnosis of acute and chronic inflammatory diseases, in addition to contributing to the understanding of the role of cytokines in these diseases.

## Introduction

Inflammatory diseases, including neurodegenerative diseases, autoimmune diseases, cardiovascular disease, cancer, among others, are characterized by chronic inflammation which promotes disease progression.^1, 2^ Inflammatory cytokines are key initiators and drivers of these diseases. Cytokines such as interleukin-6 (IL-6), interleukin-1 beta (IL-1β), interleukin-8 (IL-8), and tumor necrosis factor-alpha (TNF-α) are known therapeutic targets and biomarkers for these inflammatory diseases.^3-5^ Platforms for cytokine detection are therefore desirable tools for both research and clinical applications.

Cytokines are small, secreted proteins (< 40 kDa) which are produced by many cell types to regulate and influence immune response.^2^ They modulate both acute and chronic inflammation via a complex network of autocrine, paracrine, and endocrine interactions. Various cell populations can produce the same cytokine and the effects of each cytokine are pleiotropic. Different cytokines may have the same impact, suggesting redundancy, or cause a synergistic effect.^4, 6, 7^ Cytokines can serve as predictive, diagnostic, or prognostic biomarkers for inflammatory diseases. However, their precise role in the disease is not always clearly defined. In complex disease states, it can be unclear whether dysregulated cytokine signaling is a cause or result of disease, or both. The heterogeneity of inflammatory processes in disease states presents further challenges. For example, various studies of Alzheimer’s disease have shown increased, decreased, and unchanged levels of the same cytokine in cerebrospinal fluid.^8, 9^ These are relevant unanswered questions for researchers investigating inflammatory disease mechanisms and responses to therapeutics.

Conventional methods for cytokine detection, such as immunoassays and mass spectrometry, require expert personnel, expensive equipment, and long incubation periods. These techniques are therefore not suitable for continuous monitoring or for rapid point-of-care diagnosis.^10, 11^ Nanosensors are powerful detection tools which hold advantages over conventional methods. They have shown great promise in point-of-care devices, personalized medicine, and accessible techniques for early diagnostics.^12-14^ Nanosensors which utilize optical signal transduction methods are particularly attractive due to their potential for high-resolution imaging and minimally invasive *in vivo* sensing.^15^

Single-walled carbon nanotubes (SWCNT) are versatile materials that can be functionalized to detect a variety of clinically relevant molecules.^16, 17^ They emit near-infrared (NIR) fluorescence, which is ideal for biological imaging applications due to high tissue penetration depth and minimal autofluorescence of biological tissues in this region.^17-19^ SWCNT fluorescence is stable over time and not subject to photobleaching, providing substantial advantage over conventional fluorophores.^18, 20^ The unique optical properties of SWCNT are due to their structure, which can be visualized as a single sheet of graphene lattice rolled into a cylinder. The angle and diameter at which the sheet is rolled determines the chirality of the nanotube, categorized by and (n,m) index.^16, 21-23^ Each SWCNT chirality exhibits a distinct absorption and fluorescence emission band, which can be modulated by changes in the surrounding environment.^24^ SWCNT are often functionalized with a molecular recognition probe to confer analyte specificity to these fluorescence modulations.

Approaches for SWCNT-based sensor synthesis often include screening a library of synthetic polymers, computational analytics, and rational design with a molecular recognition probe.^19^ While screening approaches have enabled the discovery of new nanosensors with tunable properties, this may be time-consuming, labor-intensive, and open-ended.^25^ Computational approaches have begun to emerge with substantial benefit, however large sets of samples and data are required.^26^ Alternatively, rational sensor design often exploits previously-validated biological recognition elements with proven affinity for a specific antigen, such as an antibody or aptamer.^25, 27^

In this work, we designed a novel DNA aptamer-SWCNT optical sensor construct to detect IL-6. There have been previous studies that developed optical SWCNT-based sensors for IL-6, each with suitable sensitivity and selectivity^28, 29^. However, one such sensor relied on a rational design with a commercial antibody as the recognition element^28^, which may cause issues related to reproducibility/batch variation and large size/stability.^30-32^ The other demonstrated sensor relied on a synthetic polymer library screen, which may not be widely-available or as-yet optimized^29^. We chose to design an aptamer-based sensor for IL-6 as their synthesis is highly reproducible and cost-effective.^33-36^ We took advantage of a previously-reported aptamer with demonstrated affinity and function for IL-6.^37^ Prior SWCNT-aptamer sensors have proven to highly effective in detecting their target of interest.^33, 34, 38-40^ Those studies largely explored multi-step methods for SWCNT-aptamer complexion, though here we explored a direct one-step process wherein the aptamer both solubilizes the SWCNT and confers analyte specificity. We investigated several direct-complexation designs, as well as the kinetics, range, and stability of this sensor, and its function in complex biological media. Finally, we evaluated the sensor’s function using an in vitro disease model with activated macrophages.

## Methods

### Preparation of ssDNA-SWCNT

ssDNA sequences (Integrated DNA Technologies; Coralville, IA) which were previously obtained via in vitro selection (SELEX, or the Selective Evolution of Ligands by Exponential Enrichment) for their ability to bind to IL-6 were used to impart specificity for IL-6 in our sensor design (Table 1).^37^ The sequence 31Apt was the full-length aptamer obtained from prior studies, while 15Apt was a truncated version reported in that study. (GT)_15_ was used as a control, non-IL-6-binding ssDNA sequence of similar size as it has previously been used extensively its stable encapsulation of SWCNT. Further, we evaluated a (GT)_15_-31Apt hybrid sequence. We also used fluorescently-labeled versions (Cy3 and Cy5, respectively) of (GT)_15_ and 31Apt to evaluate their binding and stability.

**Table 1.**
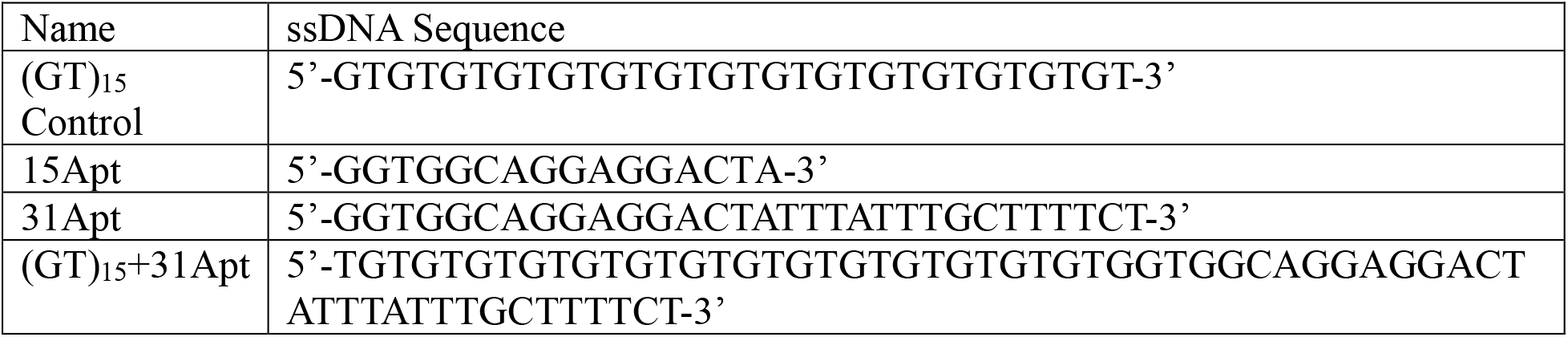
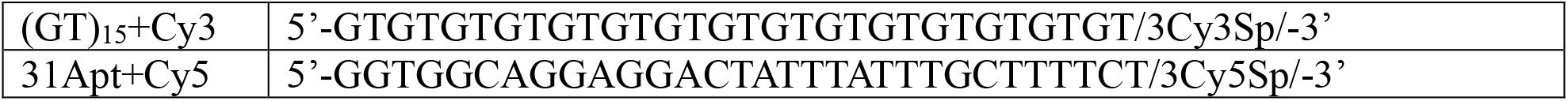
ssDNA sequences used in sensor testing.

SWCNT-ssDNA suspensions were prepared as previously described.^11, 41-44^ Briefly, HiPCO-prepared SWCNT (NanoIntegris Technologies; Boisbriand, QC) and DNA in a 1:2 mass ratio were suspended in 1X PBS. Samples were sonicated on ice at 40% amplitude for one hour by a 120 W ultrasonicator with a 1/8” microtip probe (Fisher Scientific; Hampton, NH). Sonicated suspensions were ultracentrifuged at 58,000 x g for one hour using an Optima Max-XP Ultracentrifuge (Beckman Coulter; Brea, CA). The top 75% of the suspension was collected. Samples were stored at 4°C for up to 14 days. Within 24 hours of use, samples were filtered to remove free DNA with a 100 kDa Amicon centrifugal filter (Sigma-Aldrich; St. Louis, MO) for 15 minutes at 14,000 x g. The solution retained in the filter was suspended in fresh 1X PBS.

### UV-Vis absorbance and concentration measurement

After filtration, SWCNT-ssDNA samples were subjected to absorbance measurements from 300-1100 nm using V-730 UV–vis Spectrophotometer (Jasco; Easton, MD). SWCNT-ssDNA solutions were diluted in 1X PBS to achieve absorbance values < 0.5. As in prior work,^11, 45-47^ the concentration of SWCNT was calculated using the absorbance value at the local minimum near 630 nm (extinction coefficient = 0.036 L/mg*cm)

### Near-infrared fluorescence emission measurement

NIR fluorescence spectra of SWCNT-ssDNA were primarily acquired from 900-1600 nm with a NS MiniTracer spectrophotometer (Applied NanoFluorescence; Houston, TX). The 50 mW laser source had an excitation wavelength of 638 nm. Discrete NIR fluorescence spectra were also acquired with a custom ClaIR plate reader (Photon etc; Montreal, QC) with laser source excitation wavelengths of 655 nm and 730 nm (power of ∼1750 mW). Continuous fluorescence spectra were acquired with a custom IRina probe (Photon Etc; Montreal, QC). The laser source had an excitation wavelength of 655 nm.

SWCNT (7,5), (7,6), and (9,5) chiralities were analyzed with a custom MATLAB code which fit each emission peak with a Voigt model to obtain center wavelength and maximum intensity values.^10, 11, 42^ R^2^ values for all fits were at least 0.98.

### Evaluation of SWCNT-aptamer sensitivity to IL-6

SWCNT-ssDNA of four sequences were initially screened for sensitivity to IL-6: 15Apt, 31Apt, (GT)_15_, and (GT)_15_+31Apt. SWCNT-ssDNA were tested in both buffer conditions (0.5 mg/L SWCNT in 1X PBS) and serum conditions (0.5 mg/L SWCNT in 1X PBS + 10% FBS). Baseline fluorescence measurements were taken with the MiniTracer as described above. After baseline measurements, 5250 ng/mL human recombinant IL-6 protein (Invitrogen; Waltham, MA) was added to the experimental group and an equal volume of 1X PBS was added to the control group. All samples had a total volume of 120 μL and were performed in triplicate. Fluorescence spectra were acquired with the MiniTracer at 2, 15, 30, 60, 90, 120, 150, and 180 minutes after antigen addition. Data was processed in MATLAB as previously described.

### Evaluation of sensor sensitivity and dynamic range

Due to maximal sensitivity with the 31Apt sequence, further testing was performed with this construct. 0.5 mg/L SWCNT-31Apt was tested against various concentrations of IL-6 protein in buffer conditions in triplicate: 2.1, 21, 105, 210, 2100, 5250, and 8400 ng/mL. Fluorescence spectra were acquired with the ClaIR plate reader every 15 minutes for 3 hours following antigen addition.

### Evaluation of sensor specificity

In addition to IL-6, the nanosensor was also tested against 5250 ng/mL IL-1β and 5250 ng/mL TNF-α in buffer conditions in triplicate. Fluorescence spectra were acquired with the MiniTracer as previously described.

### Kinetic response

To evaluate the kinetics of SWCNT-31Apt response to IL-6, continuous fluorescence spectra were acquired with the IRina probe. Baseline spectra were continually acquired for 300 seconds, 5250 ng/mL IL-6 was added to 0.5 mg/L SWCNT-31Apt, then spectra were continually acquired for an additional 900 seconds.

### Studies to understand the SWCNT-aptamer sensing mechanism

#### Thermal denaturation and refolding of DNA aptamer construct

SWCNT-31Apt constructs were heated to 95°C in a Thermomixer F1.5 (Eppendorf; Hamburg, Germany) for 5 minutes, then cooled on the bench until they reached room temperature (∼15 minutes) to induce reversible thermal denaturation and refolding of the aptamer. 0.5 mg/L of the heat-treated SWCNT-31Apt constructs were tested against 5250 ng/mL IL-6 in buffer conditions and fluorescence spectra were acquired in triplicate with the MiniTracer as previously described.

#### Surface passivation with BSA

Prior work has shown that surface passivation of SWCNT with BSA can decrease nonspecific interactions of nanosensors in serum via adsorption to exposed regions of the nanotube.^41, 42^ SWCNT-31Apt was incubated with BSA for 30 minutes at a 1:50 mass ratio on ice. The passivated nanosensors were then tested in triplicate against 5250 ng/mL IL-6 in serum conditions. Fluorescence spectra were acquired with the MiniTracer as previously described.

#### Noncovalent functionalization with SDBS

Prior work has shown SDBS-mediated enhancement of SWCNT-ssDNA sensor response via surface adsorption to exposed regions of the nanotube surface.^11^ 0.5 mg/L SWCNT-31Apt was tested against 5250 ng/mL IL-6 in 1X PBS with the addition of 0.2% SDBS. Fluorescence spectra were acquired in triplicate with the MiniTracer as previously described.

### Baseline sensor stability

To assess the baseline stability and shelf life of the nanosensors, a batch of SWCNT-31Apt was prepared as above, excluding the step of filtration, and stored at 4°C for 14 days. On days 1, 2, 3, 7, 8, 9, 12, 13, and 14 following suspension preparation, a 30 μL aliquot was removed from the stock solution, filtered with a 100kDa Amicon centrifugal filter as outlined above, and diluted to 0.5 mg/L in 1X PBS. Immediately following dilution, fluorescence spectra of SWCNT-31Apt were acquired in triplicate every 15 minutes for 3 hours using the ClaIR plate reader.

### DNA displacement study

To further understand the mechanism of the SWCNT-31Apt sensor and evaluate the stability of the SWCNT/aptamer/antigen complexes, a DNA displacement study was performed using cyanine dye-functionalized ssDNA. SWCNT-ssDNA suspensions were prepared as previously described using (GT)_15_+Cy3 or 31Apt+Cy5. Each SWCNT-ssDNA construct was incubated overnight at 4°C under one of three conditions: 1X PBS, 1X PBS + 5250 ng/mL IL-6 protein, or 1X PBS + 2.5% DOC, which is a positive control for displacing DNA from SWCNT. After 18 hours of incubation, all samples were subjected to centrifugal filtration with a 100 kDa Amicon centrifugal filter. The SWCNTs remaining inside the filter were discarded and the flow-through was collected for further analysis. The visible absorbance of all solutions was measured with a Biotek microplate reader with Gen5 software (BioTek; Winooski, VT). Data was analyzed in Microsoft Excel.

### Investigation of sensor functionality in an activated macrophage disease model

The RAW 264.7 murine macrophage cell line (American Type Culture Collection; Gaithersburg, MD) was seeded in 6-well plates at 100,000 cells/well. Cells were cultured with DMEM supplemented with 10% FBS and 1% penicillin-streptomycin until they reached 80% confluency. Each well was treated with 2.5 μg/mL LPS isolated from E. Coli 0111:B4 (MilliporeSigma; Burlington, MA) in 1.3 mL total volume. Cells were then incubated for 24 hours to induce macrophage activation and cytokine release.^48^ ELISA (Invitrogen, Waltham, MA, Product #88-7066-22) was performed to quantify the level of IL-6 present in the cell media. Conditioned media were collected and frozen at -20°C until use. 0.5 mg/L SWCNT-31Apt was added to thawed conditioned media samples from 0 μg/mL and 2.5 μg/mL LPS treatment groups as well as fresh DMEM + FBS as a control. Samples were evaluated in triplicated and fluorescence spectra were acquired with the MiniTracer as described above.

### Data analysis

NIR fluorescence corresponding to individual nanotube chirality emission peaks were fit with a custom MATLAB code to a Voigt model to determine their center wavelength and maximum intensity values (MATLAB code is available upon request). Changes in SWCNT center wavelength and maximum intensity values were reported relative to their emission prior to antigen addition. Statistical significance was determined with a two-sample t-test with Welch correction.

## Results and Discussion

### Screening of optimal aptamer sensor format

SWCNT separately encapsulated with four ssDNA sequences were prepared and tested against 5250 ng/mL IL-6 in buffer conditions: (GT)_15_, (GT)_15_+31Apt, 31Apt, and 15Apt (**Table 1**). Optical characterization of nanosensor constructs showed effective suspension of SWCNT with ssDNA with bright, stable NIR fluorescence emission and excitation peaks (**Figures 1A and 1B**). In prior work, evaluation of fluorescence response of 0.5 mg/mL nanosensor to 5250 ng/mL IL-6 was suitable to confirm basic nanosensor functionality.^10^ Analysis of the (7,5) SWCNT chirality showed that all control groups (containing only SWCNT-ssDNA in 1X PBS) retained a relatively stable fluorescence intensity over three hours (100% normalized fluorescence). As expected, (GT)_15_ showed no response to IL-6 protein (p = 0.65903), as it was used as a control sequence which binds to SWCNT well but with no known affinity for IL-6 (**Figure 1C**). The truncated 15Apt sequence demonstrated minimal response, quenching 15% (p = 0.00287). The fluorescence of (GT)_15_+31Apt quenched 32% (p = 0.00171), while 31Apt showed the most significant response, quenching 48% over three hours (p = 0.000496). SWCNT-31Apt exhibited time-dependent quenching over three hours, with the majority of the response occurring in the first hour (**Figure 1D**). The other three SWCNT-ssDNA construct demonstrated similar time-dependence relative to their total response (**Figure S1)**.

**Figure 1.**
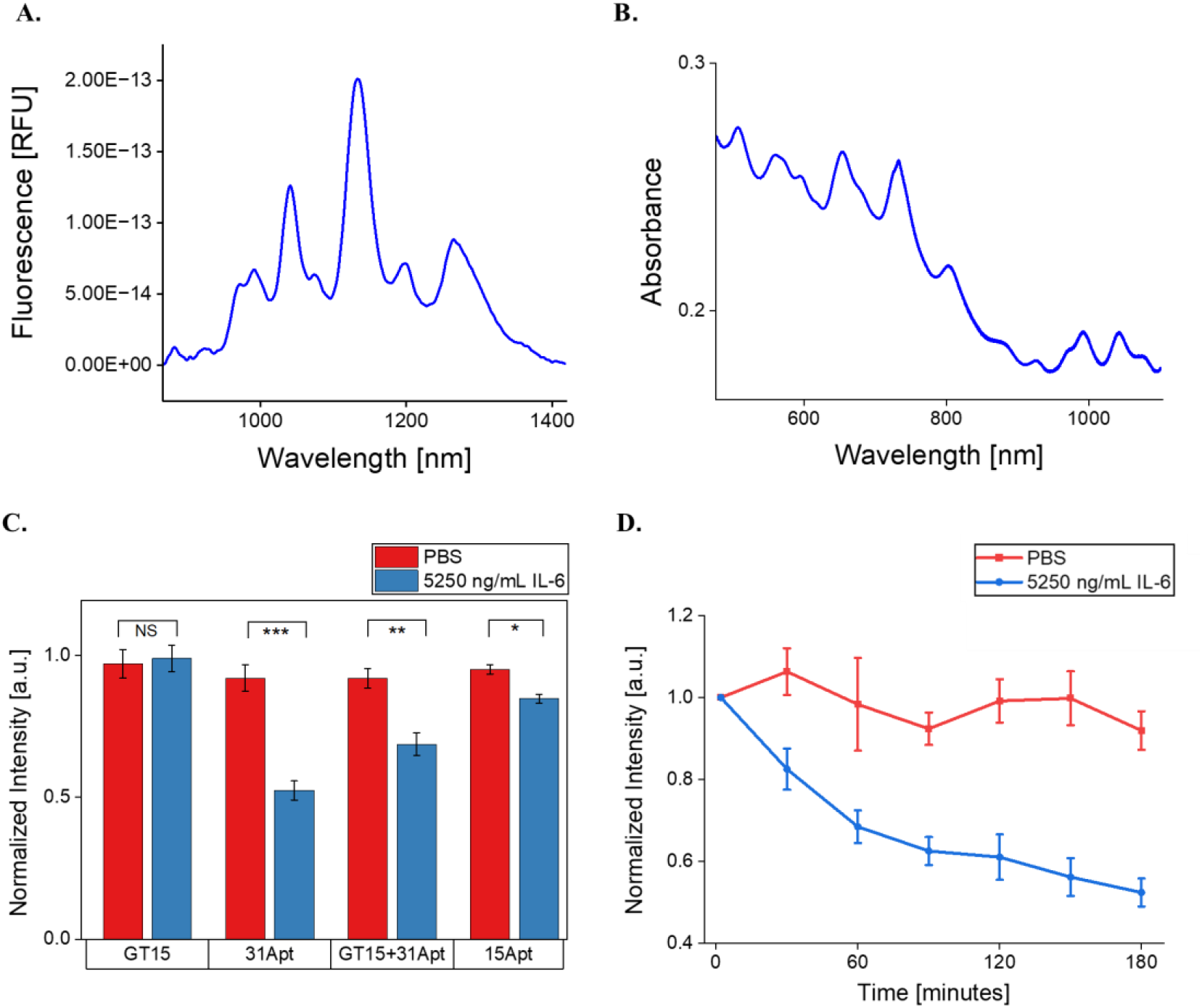
Evaluation of sensor synthesis and response to IL-6 in PBS. A) Representative NIR fluorescence spectrum of the SWCNT-31Apt construct. B) Representative Vis-IR absorbance spectrum of the SWCNT-31Apt construct. C) (7,5) Fluorescence intensities of SWCNT-ssDNA in PBS after 3 hours of exposure to IL-6. (GT)_15_: p = 0.65903, 31Apt: p = 0.000496, (GT)_15_+31Apt: p = 0.00171, 15Apt: p = 0.00287. D) (7,5) Fluorescence intensity of SWCNT-31Apt in response to IL-6 over 3 hours.

Dispersion of SWCNT with various proteins, synthetic polymers, surfactants, or oligonucleotides is an excellent way to solubilize the nanotubes and potentially impart increased biocompatibility and molecular sensitivity. However, this technique typically creates a trade-off between molecular specificity and quantum yield of the resulting nanosensors. SWCNT dispersed in surfactants exhibit high quantum yield but have no inherent selectivity towards any particular analyte.^25, 49^ Protein-wrapped SWCNT have demonstrated successful differentiation between molecularly similar targets. However, protein wrapping is limited by low dispersion efficiency and lack of concise control during protein immobilization, which can lead to unfavorable conformations for antigen binding.^50, 51^ DNA is one of the most well-studied wrappings for SWCNT sensors due to its ability for high dispersion efficiencies and customization of properties with various oligonucleotide sequences and structures. However, many SWCNT-DNA sensors do not exhibit the inherent high specificity and selectivity of their protein-based counterparts. DNA aptamers hold a unique advantage in this context advantage: as oligonucleotides, they enable high-yield SWCNT dispersions. Due to their unique structural properties, they can also provide high specificity towards a particular target.

15Apt and 31Apt were previously selected for their affinity for IL-6 via SELEX. 31Apt has also been validated for use in electrochemical and field-effect transistor sensors.^52-55^ (GT)_15_ is known to stably disperse SWCNT but does not have demonstrated affinity for IL-6.^56^ Previous aptamer-based SWCNT sensors have utilized aptamer-anchor sequences to enable ideal conformation of aptamer binding site and enhanced protein recognition.^38^ Using this rationale, we included GT15+31Apt in our study. However, the (GT)_15_-aptamer hybrid sequence was not more effective in sensing IL-6, as 31Apt alone showed the best sensitivity to IL-6. This implies that attachment of 31Apt to the nanotube surface does not inhibit the accessibility of IL-6 to its active site, and further suggests that interaction of the aptamer with the SWCNT surface is itself important in the mechanism of sensor function.

The same four SWCNT-ssDNA constructs were tested under 10% serum conditions to investigate their function in more complex protein environments. The fluorescence of the (7,5) SWCNT chirality dispersed with 31Apt quenched 32% in response to 5250 ng/mL IL-6 protein (p = 0.000624) (**Figure 2A**). The other ssDNA sequences tested showed no significant response ((GT)_15_: p = 0.94660, (GT)_15_+31Apt: p = 0.58306, 15Apt: p = 0.1526). Quenching of 31Apt reached its maximum within 30 minutes, after which it was stable (**Figure 2B**). No other SWCNT-ssDNA construct demonstrated any substantial differences across three hours (**Figure S2)**. 31Apt was therefore used in further sensor testing.

**Figure 2.**
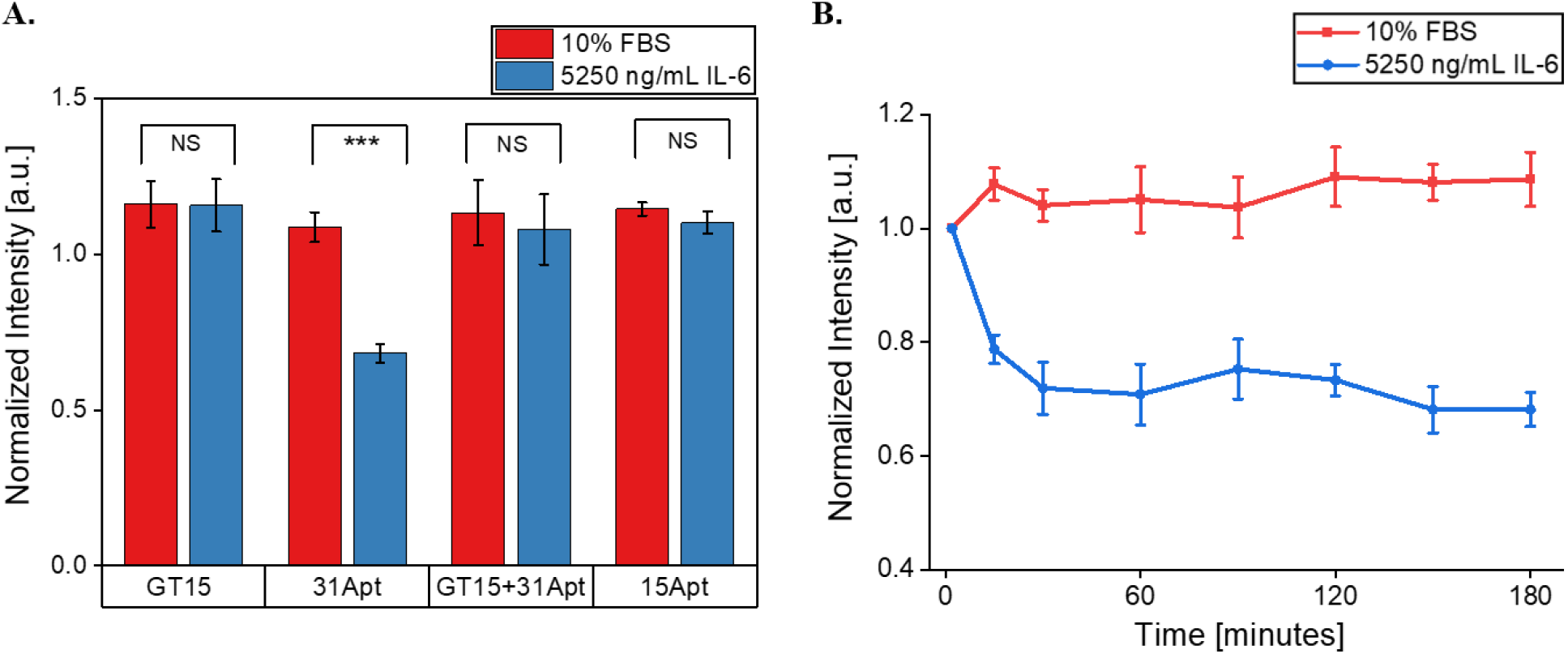
Sensor response to IL-6 in PBS + 10% FBS. A) (7,5) chirality fluorescence intensities of SWCNT-ssDNA after 3 hours of exposure to IL-6 + 10% FBS. (GT)_15_: p = 0.94660, 31Apt: p = 0.000624, (GT)_15_+31Apt: p = 0.58306, 15Apt: p = 0.1526. B) (7,5) chirality fluorescence intensity of SWCNT-31Apt over 3 hours in response to 5250 ng/mL IL-6 + 10% FBS.

### Sensitivity and dynamic range of the SWCNT-31Apt sensor

We next evaluated the sensitivity of the SWCNT-31Apt sensor to respond to a range of IL-6 concentrations in PBS. At or below 21 ng/mL IL-6, SWCNT-31Apt did not show any significant quenching of photoluminescence (2.1 ng/mL: p = 0.06744, 21 ng/mL: p = 0.42251) (**Figure 3A**). At or above 105 ng/mL IL-6, SWCNT-31Apt demonstrated quenching of fluorescence intensity that was monotonic to the sample concentration of IL-6 (105 ng/mL: p = 0.02783, 210 ng/mL: p = 0.00216, 2100 ng/mL: p = 0.000708, 5250 ng/mL: p = 0.00777, and 8400 ng/mL: p = 0.000231). The highest concentration tested, 8400 ng/mL, induced the most dramatic response (58% fluorescence quenching). We therefore conclude this sensor construct is quantitative. Pathological levels of IL-6 in biofluids span 0.005-500 ng/mL, depending on the fluid and disease.^57^ The nanosensor demonstrated monotonic response between 105-8400 ng/mL, with a limit of detection within the clinically relevant range.

**Figure 3.**
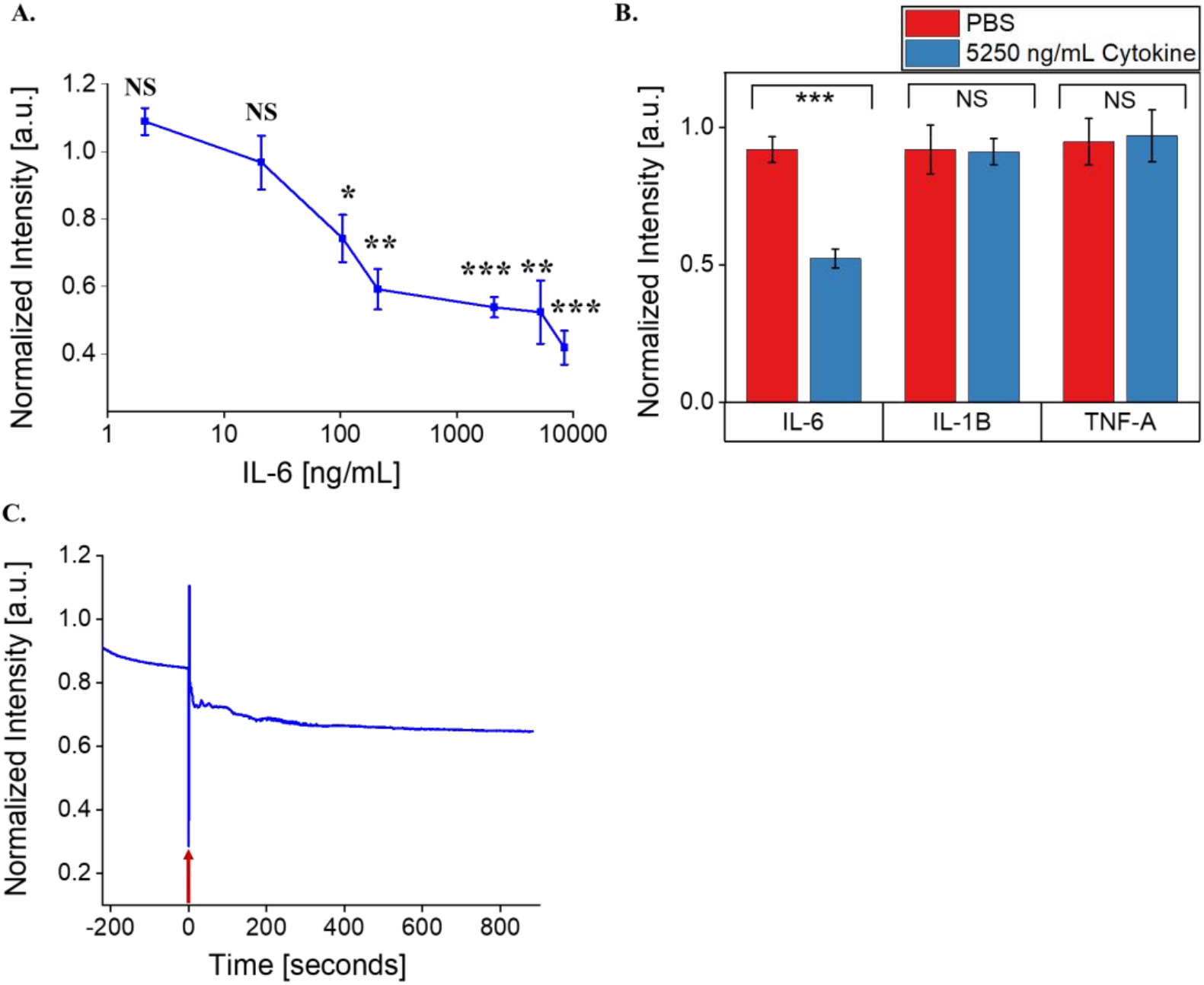
Determination of nanosensor sensitivity, selectivity, and rapidity. A) (7,5) Fluorescence response of SWCNT-31Apt in PBS as a function of IL-6 concentration. B) (7,5) Fluorescence intensity of SWCNT-31Apt in PBS after 3 hours of exposure to 5250 ng/mL IL-6, IL-1β, and TNF-α. C) Fluorescence intensity of SWCNT-31Apt in PBS immediately before and after the addition of 5250 ng/mL IL-6. Moment of protein addition (time 0) indicated by red arrow.

### Specificity of the SWCNT-31Apt sensor

In order to determine if sensor response is specific to IL-6 and not other inflammatory cytokines in an inflammatory disease context, we tested its response against two other cytokines. When incubated with inflammatory cytokines IL-1β and TNF-α, the sensor exhibited no significant fluorescence modulation (**Figure 3B**). IL-1β and TNF-α were chosen for this application due to their structural and immunological similarities to IL-6.^2, 58^ All three are considered classical pro-inflammatory cytokines known to be dysregulated in inflammatory disease states.^4^

### Rapid kinetic response

Continuous acquisitions of fluorescence spectra of SWCNT-31Apt showed that quenching of photoluminescence begins within seconds of IL-6 addition (**Figure 3C**). 25% fluorescence quenching occurred within 15 seconds of antigen addition. Fluorescence intensity steadily decreased: 30% quenching within 115 seconds and 35% quenching within 560 seconds of antigen addition. This finding is consistent with previous time-dependent response of the sensor (**Figures 1D and 2B**) where at least 20% quenching occurs within 15 minutes of antigen addition and quenching continues steadily over the course of three hours. Detection and quantification of biomarkers with temporal resolution is beneficial for clinical and research applications. Due to its ability to respond rapidly to IL-6, this nanosensor construct could be used in immunology research to study the kinetics associated with cytokine release in inflammatory disease states. Similarly, dynamic and time-sensitive measurements of biomarkers are ideal for clinical applications that require continuous monitoring.

### Investigation of sensing mechanism

We performed several modifications to the standard assays above to further understand the mechanism by which the SWCNT-aptamer complex responds with high sensitivity, specificity, and speed to IL-6. Typically, in DNA aptamer selection experiments, the library is heated to 95°C and cooled back to room temperature to ensure proper tertiary structure is adopted. We therefore tested whether denaturation and folding affected sensor function. However, after heating to 95°C and cooling back to room temperature, the SWCNT-31Apt construct no longer exhibited the same response to IL-6. Rather, it showed only 10% quenching after 3 hours (p = 0.03216) (**Figure S3A)**. This suggests that the probe-tip sonication of aptamer with SWCNT is sufficient for proper folding, and that additional perturbation of this interaction disrupts the ability to respond to IL-6.

Surface adsorption of bovine serum albumin (BSA) has been shown to enhance the functionality of an antibody-based SWCNT sensor through adsorption to exposed surfaces of the nanotube.^41, 42, 59^ We therefore evaluated whether such adsorption to exposed SWCNT surface would perturb the sensor response to IL-6. In this study, BSA passivation did not enhance the selectivity of SWCNT-31Apt when deployed in serum conditions (**Figure S3B**). In fact, there was no significant difference in the fluorescence intensity of BSA-incubated SWCNT-31Apt with and without the presence of IL-6 in serum conditions (p = 0.44531). We therefore conclude that this study supports our hypothesis that the SWCNT interaction with the 31Apt sequence is essential to IL-6 sensing, as perturbation of that interaction with a large BSA molecule (66 kDa compared to ∼19 kDa of the aptamer) interferences with such a response.

Similarly, additional prior work has investigated the interaction of surfactants with the nanotube surface.^49^ As an example, a previous SWCNT sensor for microRNA was substantially enhanced due to the interaction of the hydrophobic surfactant sodium dodecylbenzene sulfate (SDBS) with exposed SWCNT surface and replacing ssDNA removed from the SWCNT surface.^11^ In this study, however, SWCNT-31Apt showed no significant response to IL-6 in the presence of 0.02% SDBS (p = 0.08707) (**Figure S3C)**. We believe this result further supports our hypothesis that 31Apt is not removed from the SWCNT surface and that the 31Apt sequence must be free to interact or potentially rearrange on the nanotube surface to exhibit detection of IL-6.

### Stability of the sensor

To further investigate the mechanism of sensor function, and to understand its stability, we first investigated its stability over the course of 14 days. To do so, we measured the baseline fluorescence of SWCNT-31Apt, finding considerable stability.

Fluctuations of all chiralities analyzed –the (8,3), (7,5), and (7,6)—were minimal and within 10% variation (**Figure 4A)**. This suggests that the SWCNT-31Apt construct is stable for at least two weeks when stored at 4°C.

**Figure 4.**
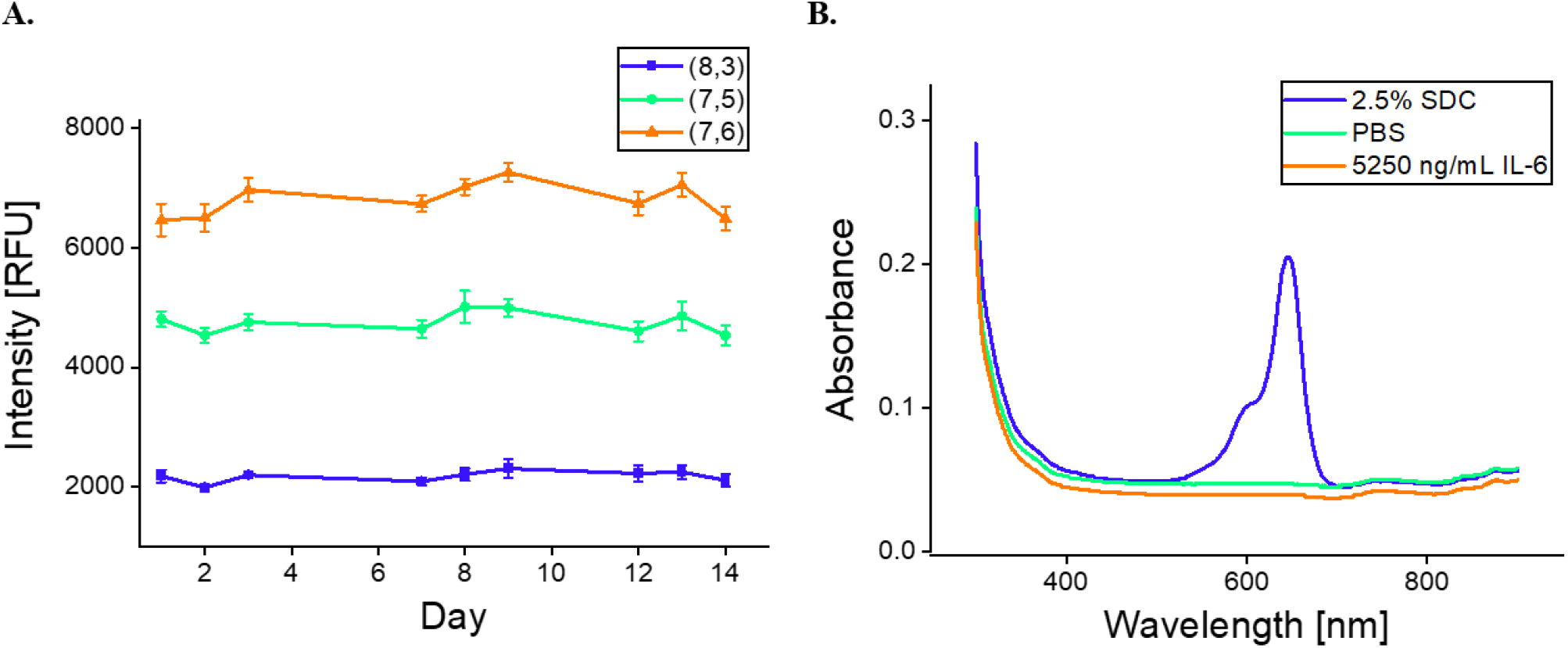
Stability of nanosensor constructs. A) (8,3), (7,5), and (7,6) Fluorescence intensity of SWCNT-31Apt in PBS for 2 weeks after preparation. B) Absorbance of flow-through from SWCNT-31Apt+Cy5 after overnight incubation.

We next studied whether the interaction of 31Apt with IL-6 displaces the aptamer sequence from the surface of the nanotube. One possibility for the mechanism of sensing was that the sensor quenching response may be due to such a displacement and subsequent SWCNT aggregation. In this experiment, the results suggested that IL-6 protein does not displace 31Apt from the SWCNT surface during sensor response. When SWCNT functionalized with cyanine dye-labeled ssDNA ((GT)_15_+Cy3 or 31Apt+Cy5) was incubated overnight with 2.5% of the surfactant deoxycholate (DOC) as a positive control, DNA was displaced from the nanotube surface as evidenced by fluorescence flow-through of the centrifugal filter membrane (**Figure S4)**. The same nanotube constructs incubated with PBS as a negative control demonstrated no DNA displacement. SWCNT-31Apt+Cy5 saw no measurable DNA displacement after overnight incubation with 5250 ng/mL IL-6 protein (**Figure 4B**). This is promising evidence for future applications of the sensor, where long-term storage and stable binding would be beneficial, as well as the potential for sensor reversibility. Importantly, this result demonstrates that the IL-6 interaction with the aptamer occurs on the surface of the nanotube and not independent of it. This further suggests that the 31Apt may undergo rearrangement on the SWCNT surface to allow IL-6 binding and fluorescence response but does not undergo dissociation or release.

### Investigation of sensor functionality in activated macrophages

To evaluate the function of the SWCNT-31Apt nanosensor construct using an in vitro model of disease, in this case acute bacterial infection, we exposed the sensor to conditioned cell media. The sensor was exposed to non-conditioned media as a baseline control, media recovered with naïve RAW-264.7 murine macrophages, or media recovered from the same macrophages activated with bacterial lipopolysaccharides (LPS). By ELISA, we found that macrophages treated with 2.5 μg/mL LPS for 24 hours expressed 4135.9 pg/mL IL-6 in their conditioned media, while macrophages not treated with LPS expressed 0.8 pg/mL IL-6 in their conditioned media. The nanosensor demonstrated was a 22% decrease in fluorescence from the non-LPS conditioned media to the LPS conditioned media (p = 0.0014) (**Figure 5A**). The observed response aligns with our previously constructed concentration-response curve, as 20% quenching would be expected at the given concentration of IL-6 (**Figure 3A**). As previously observed, the majority of the sensor response was observed in the first 15 minutes then continued steadily over three hours (**Figure S5A)**.

**Figure 5:**
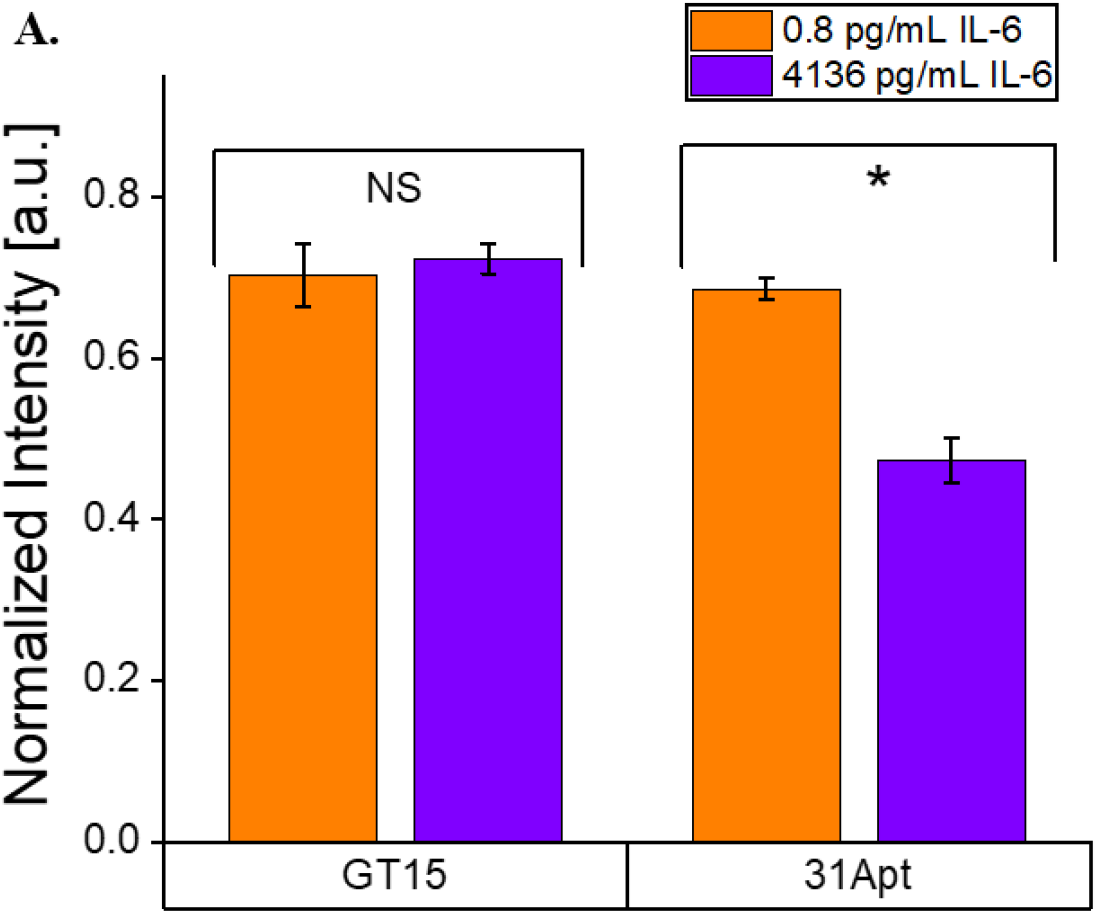
Quantification of IL-6 in conditioned media from LPS-activated macrophages. A) (7,5) fluorescence intensity of SWCNT-(GT)_15_ and SWCNT-31Apt after 3 hours of exposure to conditioned media from Raw 264.7 cells. SWCNT-31Apt distinguished activated and non-activated macrophages (p = 0.0014) while SWCNT-(GT)_15_ did not (p = 0.48275).

To determine the role of 31Apt in the observed sensor response, we also performed the experiment with SWCNT-(GT)_15_. This non-sensing control exhibited no significant difference in the non-LPS conditioned media and LPS conditioned media (p = 0.48275) (**Figure 5B, Figure S5B**), demonstrating the importance of the IL-6-specific aptamer in sensor functionality. As we previously found that this nanosensor is highly specific to IL-6, having demonstrated a consistent response in 10% FBS and no response to other cytokines, we conclude that this response is IL-6 specific in conditioned media despite the presence of other cytokines.^48^ We were also excited to find that the sensor responds as well to full-length mouse IL-6 as it did recombinant human IL-6 in the above studies.

## Conclusions

In this work, we designed and investigated the use of a novel IL-6 aptamer-based sensor complexed with SWCNT as an optical transducer. Sensor construction was straightforward with no intermediary steps, and we found the sensor was most optimally functional with no hybridization sequence or truncation. We also found that the sensor response requires the IL-6 aptamer and not a random ssDNA sequence. We further investigated the sensitivity, selectivity, rapidity, and mechanism of action/stability of the nanosensor. The sensor demonstrated a clinically relevant detection limit of 105 ng/mL and responded specifically to IL-6 in buffer and serum conditions.^57^ This nanosensor construct exhibited rapid response to its target and forms a stable construct with IL-6 without displacing the aptamer from the SWCNT surface. Excitingly, we demonstrated that this nanosensor can detect bacterial infection, as modeled by LPS-incubated macrophages in vitro.

Ongoing work is likely to push this novel sensor further towards utility both in the clinic and as a research tool. It may be possible to develop multiplexed nanosensors that detect several cytokines with chirally-separated SWCNT, which may also improve sensitivity and specificity of the sensor.^60, 61^ It may also find utility as in prior work in SWCNT-based sensing of extracellular analytes via sensor immobilization on a glass substrate upon which cells are cultured.^62^ Further in vivo development may be possible through embedding the sensor within a semipermeable membrane or hydrogel.^63^ We anticipate that a combination of these approaches may confer clinical utility in early-stage detection of chronic inflammatory diseases or rapid detection of infection.

## Acknowledgements

The authors wish to acknowledge all members of the Williams Lab for discussion and feedback. This work was supported by NIH R35GM142833 (R. Williams) and the support of The City College of New York Grove School of Engineering. A. Ryan was supported by a G-RISE Ph.D. traineeship from the National Institutes of Health (T32GM136499).

