## Supplementary Information for "An optical aptamer-based cytokine nanosensor detects macrophage activation by bacterial toxins"

### Supplementary Figures

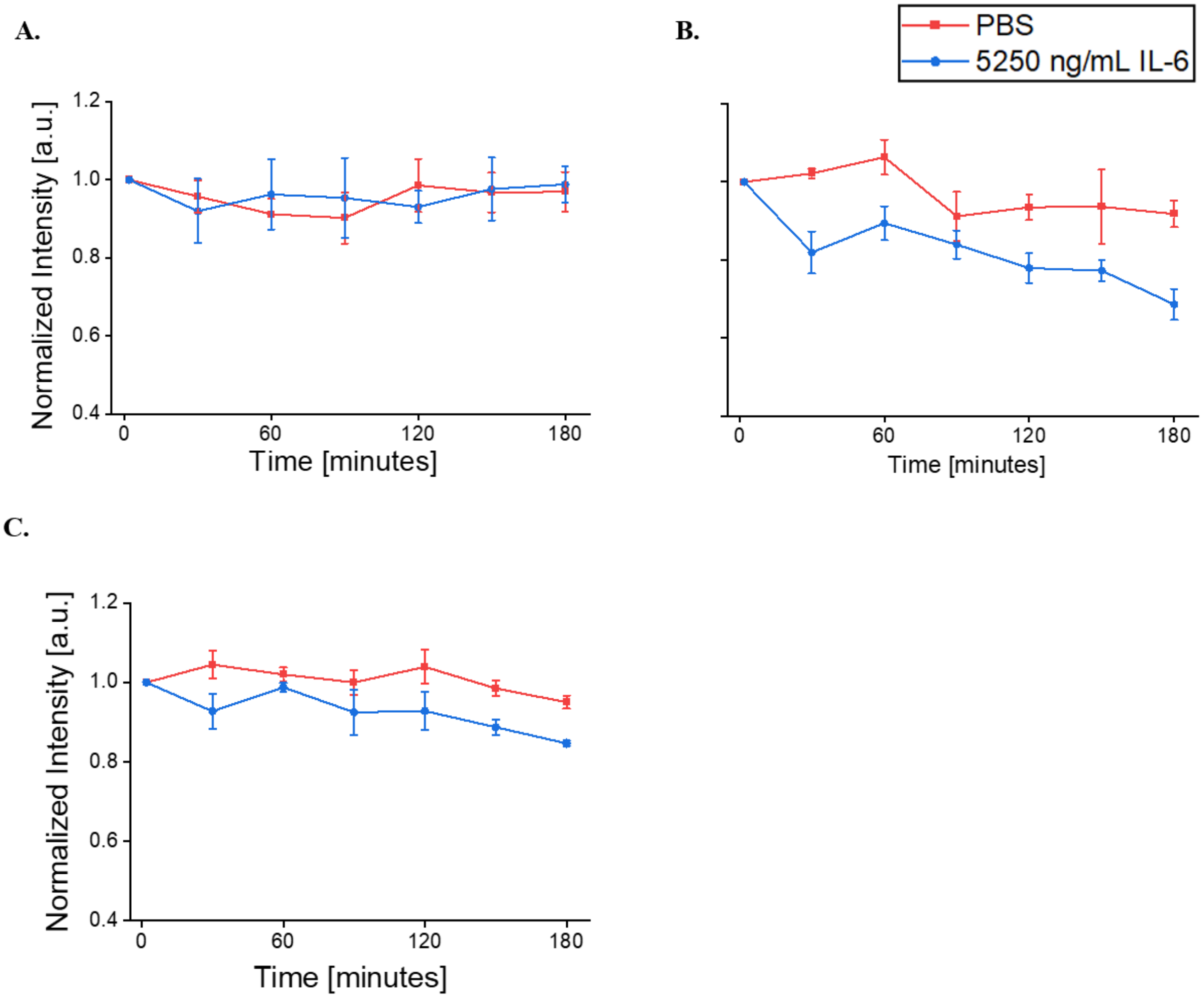


**Figure S1**. **Time-dependent response of SWCNT-ssDNA constructs to IL-6 in PBS.** A. (7,5) fluorescence intensity of SWCNT-$(GT)_{15}$ over three hours. B. (7,5) fluorescence intensity of SWCNT-$(GT)_{15}$+31Apt over three hours. C. (7,5) fluorescence intensity of SWCNT-15Apt over three hours

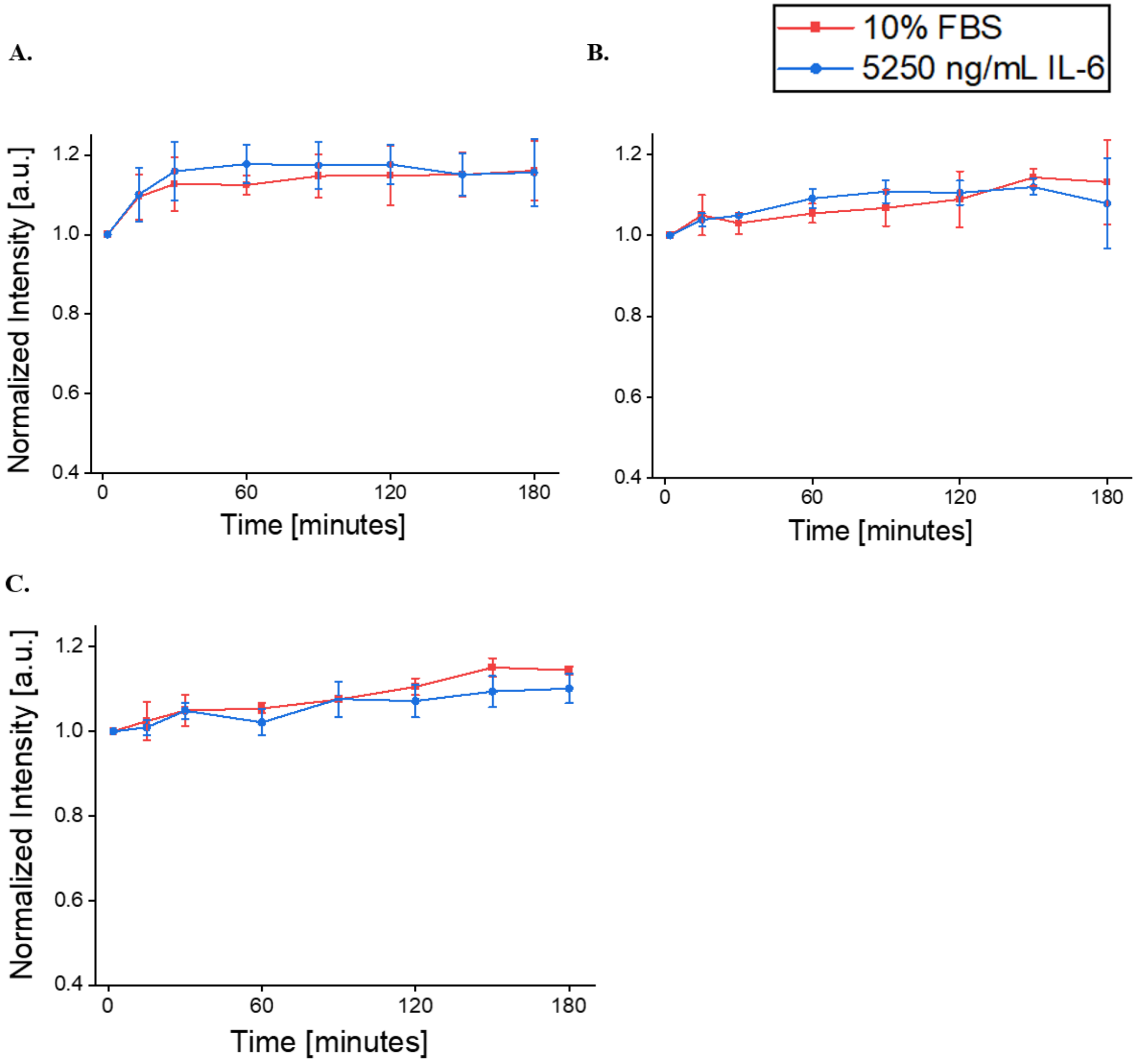


**Figure S2. Time-dependent response of SWCNT-ssDNA constructs to IL-6 in PBS + 10% FBS.** A. (7,5) fluorescence intensity of SWCNT-$(GT)_{15}$ over three hours. B. (7,5) fluorescence intensity of SWCNT-$(GT)_{15}$+31Apt over three hours. C. (7,5) fluorescence intensity of SWCNT-15Apt over three hours

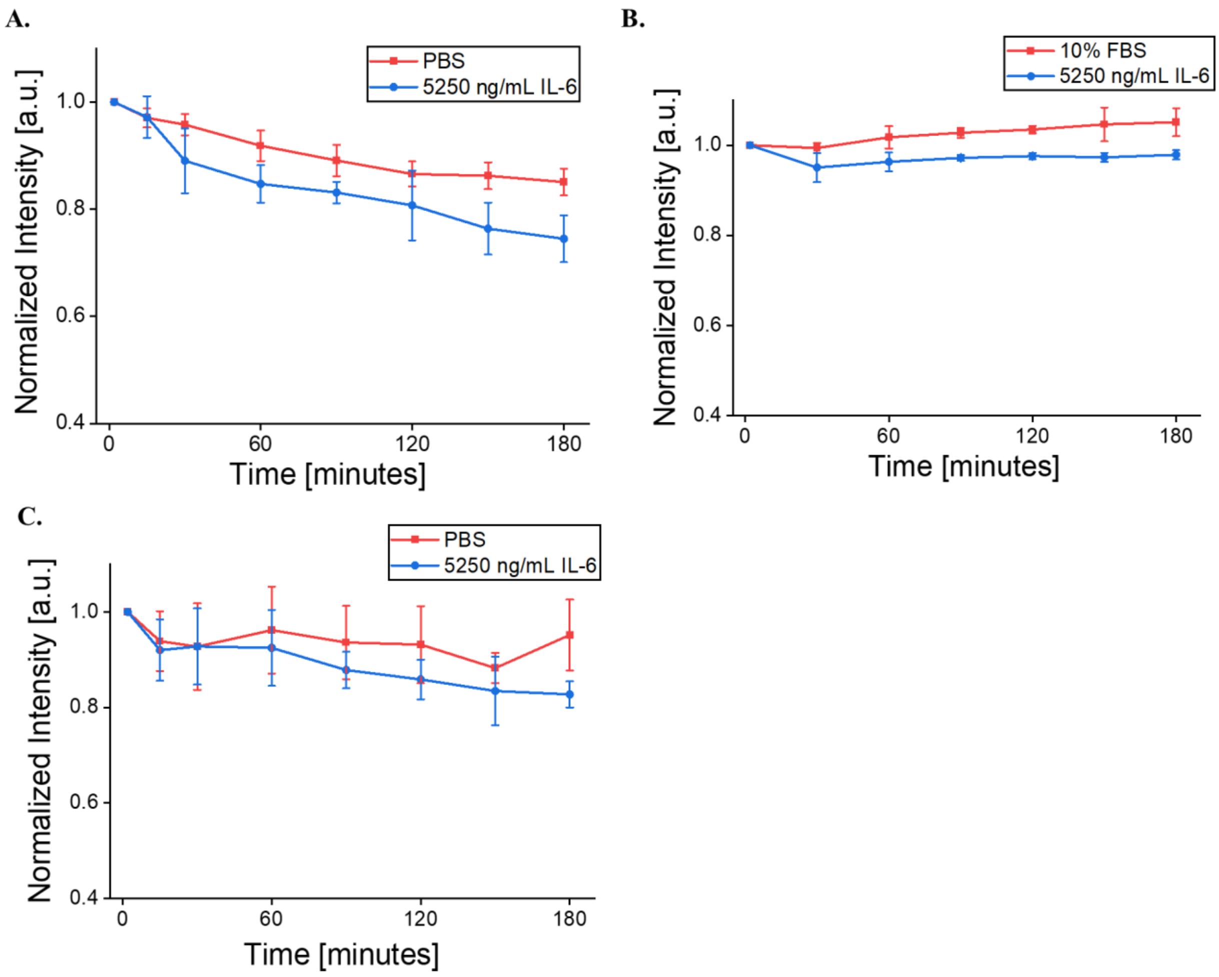


**Figure S3. Mechanistic modulations of nanosensor**. A. (7,5) Fluorescence intensity of heat-treated SWCNT-31Apt in response to IL-6 in PBS over three hours. B. (7,5) Fluorescence intensity of BSA-passivated SWCNT-31Apt in response to IL-6 in PBS + 10% FBS over three hours. C. (7,5) Fluorescence intensity of 0.02% SDBS-coated SWCNT-31Apt in response to IL-6 in PBS over three hours.

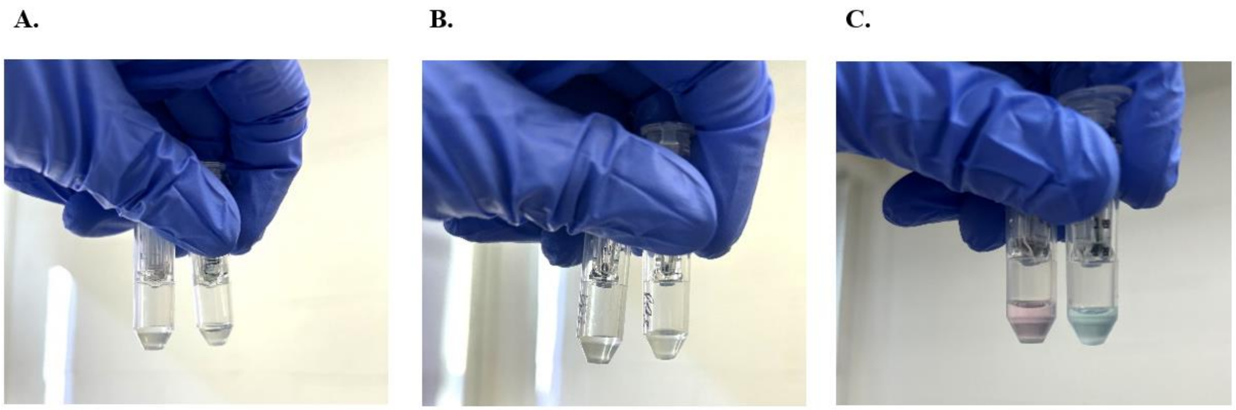


**Figure S4. Photos of displaced cyanine dye-labeled DNA**. A. Flow-through after incubation of SWCNT-$(GT)_{15}$+Cy3 (left) and SWCNT-31Apt+Cy5 (right) with PBS. B. Flow-through after incubation of SWCNT-$(GT)_{15}$+Cy3 (left) and SWCNT-31Apt+Cy5 (right) with 5250 ng/mL IL-6 protein in PBS. C. Flow-through after incubation of SWCNT-$(GT)_{15}$+Cy3 (left) and SWCNT-31Apt+Cy5 (right) with 2.5% DOC.

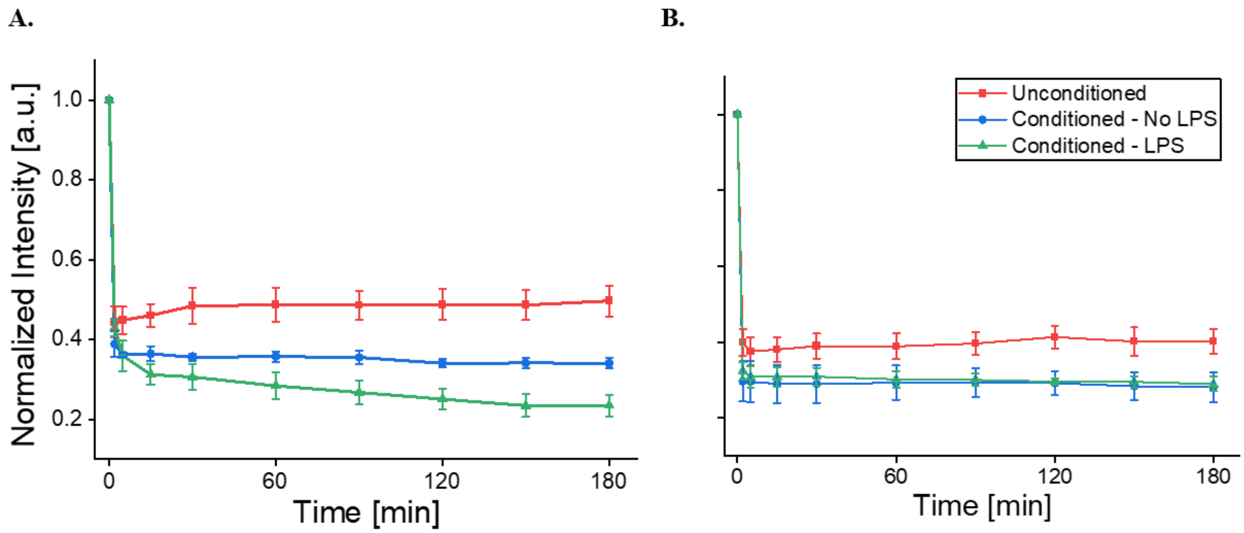


**Figure S5. Time-sensitive response of SWCNT-ssDNA to conditioned media from Raw 264.7 cells.** A) (7,5) Fluorescence intensity of SWCNT-31Apt in response to cell media samples over three hours. B) (7,5) Fluorescence intensity of SWCNT-$(GT)_{15}$ in response to cell media samples over three hours.
